# TATAT: a containerized software for generating annotated coding transcriptomes from raw RNA-seq data

**DOI:** 10.1101/2025.07.09.663867

**Authors:** Alexander J. Brown, Stephanie N. Seifert

## Abstract

**Motivation:** Many transcriptome creation workflows are not standardized, are difficult to install or share, prone to breaking as dependencies update or cease to be maintained and are resource intensive. Due to a lack of authoritative literature, many also overlook potentially important steps, such as thinning contig over-assembly or identifying transcript consensus across samples, which reduce resource demands during annotation and increase the accuracy of final transcripts.

**Results:** We developed TATAT, a modular, Dockerized software that contains all the tools necessary to generate an annotated coding transcriptome from raw RNA-seq data. The tools remain in a static state and can be coordinated with bash and python scripts provided therein, making TATAT a standardized, reproducible workflow that can easily be shared and installed. We preferentially incorporate tools that are not only accurate, but are fast and require less RAM, and subsequently show TATAT can generate a comprehensive transcriptome for a non-model organism, the Egyptian rousette bat (*Rousettus aegyptiacus*), in ∼8 hours in a high-performance computing (HPC) environment.

**Availability and implementation:** The TATAT code, instructions, and tutorial are available at https://github.com/viralemergence/tatat.

## 1 Introduction

Quantifying changes in gene expression remains a foundational approach to understanding how an organism is affected by, and responds to, a stimulus. However, many organisms of interest do not have either complete genomes or transcriptomes sequenced (1). If a researcher desires to study these non-model organisms and perform gene-dependent analyses, they must sequence the organisms themselves. While the cost of genome sequencing has dropped dramatically (2–4), sequencing only RNA is often preferential as it is cheaper and subsequent transcriptome generation has lower computational demands (5,6).

The computational process of RNA-seq transcriptome generation can be accomplished in many ways, generally following a pattern of preprocessing sequencing reads for assembly, performing *de novo* assembly, optionally thinning the assembled contigs, and finally annotating the contigs of interest. Many intuitive, open-source tools exist to perform these steps. For instance, preprocessing sequencing reads is often accomplished via tools including Trimmomatic (7), Cutadapt (8), and fastp (9). Similarly, assembly may be carried out with SOAPdenovo2 (10), Trinity (11), and rnaSPAdes (12). Thinning, if done, can potentially be addressed with CD-HIT-EST (13) or EvidentialGene (14). Lastly, annotation can be performed with HMMER3 (15), BLAST+ (16), Diamond (17), and MMseqs2 (18).

Despite the availability and accuracy of these tools, there are still hurdles to overcome. For instance, many bioinformatic tools require other tools (dependencies) to be installed before they will function, and if those dependencies are updated or cease to be supported, it may cause the initial tool to break. For instance, EvidentialGene requires the Exonerate package (19), which is no longer hosted on EMBL-EBI, and rnaSPAdes requires python3 before it can run. Additionally, installing the sheer number of dependencies may be challenging to researchers, or conflict with preexisting versions of the software in the computing environment. As an example, the transXpress workflow (20) requires installation of over 20 dependencies before running (although it does attempt to automate this process for researchers). Also, some of the primary tools themselves may cease to be supported (e.g. Trinity no longer has funding allocated towards maintenance). Furthermore, system interoperability can be constrained by operating system and environment-specific differences, which may prevent a given tool or workflow from running consistently across different computing platforms.

Even if installation and configuration are successfully navigated, coordinating these tools into a cohesive, standardized workflow can be problematic. Some more tedious obstacles include making sure the output of one tool is an acceptable format for the input of the next tool or ensuring intermediate files and directory structures are easy to interpret. More serious issues include biological considerations such as how the transcripts should be annotated. For instance, while BLAST is identified as a usable tool, when run locally it requires a BLAST database, generally reports multiple matches for a given query sequence, and the reported matches are unintuitive accession numbers. This poses questions like, which database is most relevant? Which sequence match is most biologically relevant, especially if they scored equally well? Which genes are present in the transcriptome since the accession numbers do not explicitly indicate genes? And when more than one query sequence reports as a match to the same gene, which sequence should be used? Unfortunately, there are differing opinions and a lack of literature to definitively answer these questions and create a standardized approach.

Another concern is it has been repeatedly observed that assemblers tend to produce far more contigs than there are genes (described as “over-assembly”). Some tools have been made to “thin” the excess contigs, though many workflows do not perform thinning the same way, if at all. For instance, in a paper by Lee et al. 14,796,219 initial contigs were produced by Trinity, and CD-HIT was used to reduce the number to 4,746,293 transcripts, which were then annotated to create a final transcriptome of only 24,118 genes (21). In contrast, the transXpress (20) and Trinotate (22) workflows perform no dedicated thinning steps, although both workflows can partially address this by looking for open reading frames. This is technically problematic because annotating these excess contigs (i.e. 4,722,175) dramatically increases the computational load and runtime of transcriptome generation. But it is also scientifically challenging as it once again poses questions to researchers about which process to use and how to perform it, since there are conflicting workflows.

To overcome these many challenges and improve reproducibility of scientific findings, we developed a Dockerized software called Transcriptome Assembly, Thinning, and Annotation Tool (TATAT). By Dockerizing TATAT, all dependencies are already installed and remain versioned despite updates, and the entire software package can be downloaded in a single Docker image file and run on any operating system.

Furthermore, TATAT leverages a SQLite database and several python and bash scripts to coordinate the inputs and outputs of each of the internal bioinformatic tools. After a researcher performs basic preparation for running on their computing environment, TATAT can generate a comprehensive coding transcriptome. This design *de facto* creates a standardized, reproducible workflow. The details of TATAT’s functionality and validation are described below.

## 2 Materials and methods

### 2.1 TATAT Overview

Figure 1 depicts the overall intended workflow for TATAT, where raw RNA-seq data is converted to an annotated transcriptome. Sequencing reads may first be acquired from the Sequencing Read Archive (SRA) or in-house, then the desired reads are preprocessed with fastp, *de novo* assembled into contigs with rnaSPAdes, the excess contigs are thinned with EvidentialGene and candidate CDSs identified, and finally the CDSs are annotated by performing a BLAST search and processing the results with the NCBI’s Datasets tool (23). These steps are grouped into 3 stages: Assembly, Thinning, Annotation, and are described in detail below.

**Figure 1.**
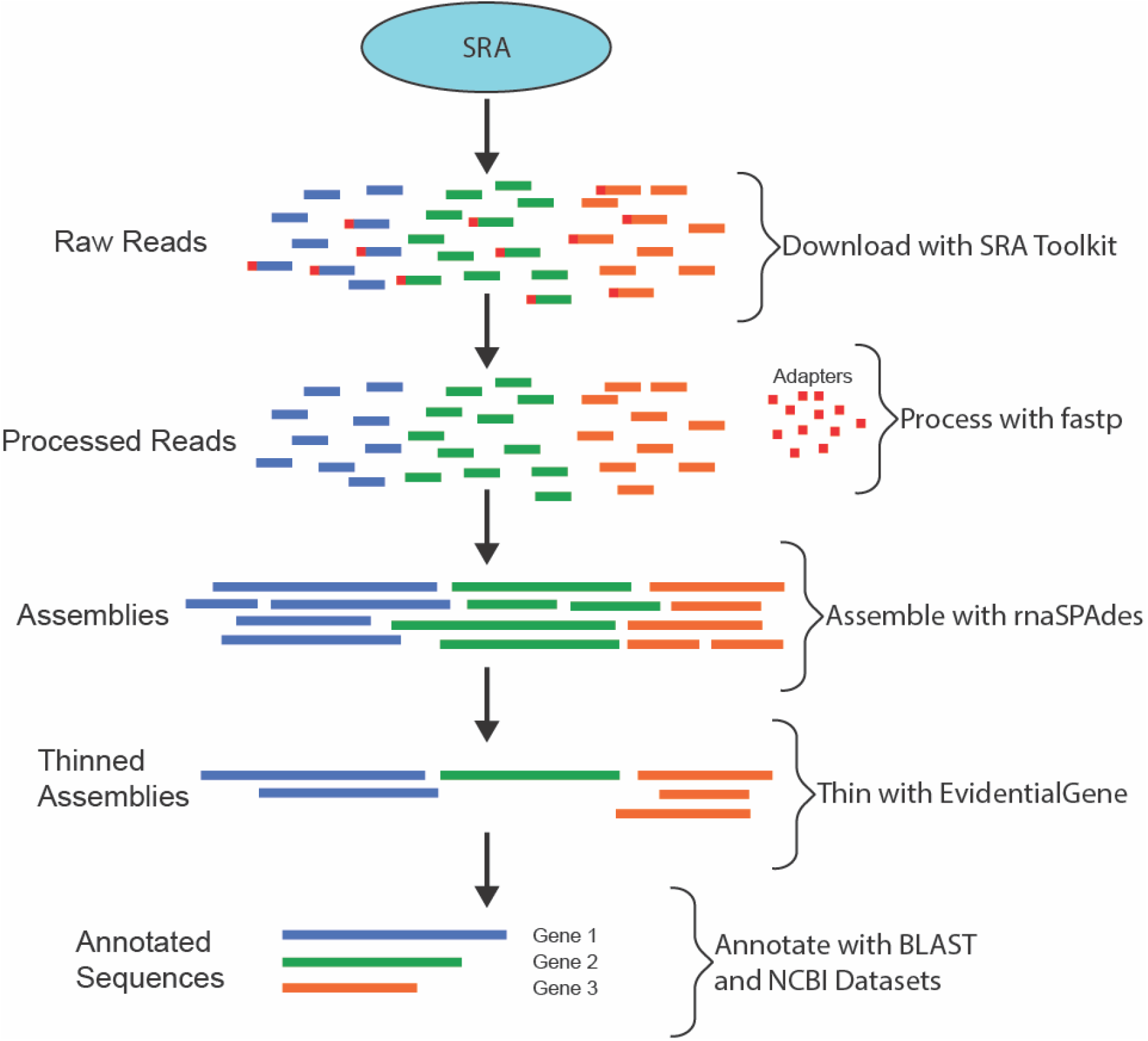
An overview of the main TATAT workflow. The TATAT workflow may start with downloading sequencing reads from the SRA. Once reads are obtained, they are processed with fastp to remove adapters and sequencing errors where possible, then *de novo* assembled into contigs with rnaSPAdes. Redundant contigs are removed and candidate CDS determined with EvidentialGene, leaving a thinned contig pool. These final candidates are annotated by leveraging BLAST and the NCBI’s tool Datasets.

#### 2.1.1 Assembly

The first (optional) step of Assembly is to acquire the RNA-seq data from the SRA using the SRA Toolkit v3.1.1. To simplify this step and all following steps, a metadata table describing the samples must be provided, which TATAT imports into a SQLite database and then uses to automatically download the sequencing data per sample, decompress the data, and move each sample’s data to a collated directory.

After this, or if the samples are already available, the reads are processed using fastp v0.26.0, which removes low quality reads, corrects for sequencing error when possible, and trims adapter sequences. The fastp tool (9) is generally faster and more accurate when compared to other sequencing read processing tools like Cutadapt and Trimmomatic, thus we leverage fastp and the default parameters for processing data. TATAT allows the user to pass in how many CPUs to use and which adapter sequences to trim. As an additional quality-of-life improvement, TATAT will automatically determine if the reads are single or paired (though not interlaced) and will modify the fastp command accordingly.

Lastly, the fastp-processed reads are *de novo* assembled into contigs using rnaSPAdes v4.0.0. We reviewed the initial publication (12) for rnaSPAdes and observed, much like fastp, it is benchmarked to be better or comparable to tools like Trinity and SOAP when comparing attributes such as accuracy, runtime, and memory usage. E.g. for the human dataset, rnaSPAdes completes in ∼5 hours and uses 32 GB RAM, whereas Trinity takes ∼18 hours and uses 50 GB RAM. Other considerations were made as well, including ease of incorporating the tools into a Docker image and announcements that some tools, namely Trinity, may not be supported in the future. In summary, we find rnaSPAdes to be both the most reliable and efficient tool for assembly and likewise use the default parameters, with the added TATAT feature to detect single or paired reads.

It is worth noting that each of these steps in the Assembly stage is intentionally decoupled so that alternative tools could be used (e.g. Cutadapt or Trinity). However, these tools are not present in the TATAT Docker image and we recommend using the default tools so that the workflow is standardized, efficient, and as minimally resource-intensive as possible.

#### 2.1.2 Thinning

Many *de novo* assembly tools produce far more transcript contigs than would be expected based on the observed or predicted number of genes in an organism. This over-assembly is the result of several factors, such as heterozygous mRNA (24), unprocessed mRNA (25), transcriptional noise (26,27), and various incorrect assemblies (25,28) (see Supplemental Figure 1). In addition, when an assembly is done per sample, each sample has the potential to produce a contig for the same transcript, meaning a common housekeeping gene like ubiquitin (UBB) might show up correctly, but redundantly, and contribute to the over-assembly.

Consequently, we choose to perform thinning of the over-assemblies to reduce runtime and resource requirements for subsequent annotation. Of the workflows that perform thinning, many take the approach of clustering sequences using tools like CD-HIT to group redundancies and fragmented transcripts, then selecting a representative sequence from the cluster (13,29). However, these tools often simply choose the longest contig to represent the cluster, even though it may be erroneous (e.g. a contig generated from DNA contamination). Likewise, these tools miss an opportunity to examine transcript consensus across samples to identify transcripts that are more likely to be real.

We identified EvidentialGene as a transcript thinning tool which takes into consideration the potential origins of contigs and looks for consensus across samples, similarity across transcripts, and identifies probable open reading frames (ORFs). Not only do papers suggest it may be more accurate than approaches like CD-HIT for choosing representative transcripts (30,31), but it also appears to more effectively thin the overall contig count.

The TATAT workflow leverages EvidentialGene (tagged 2023/07/15) to thin the contigs produced by rnaSPAdes and then identify candidate CDSs, and store that information in the SQLite database in a table called “cds”. While metadata for all CDSs are retained, the EvidentialGene tool demarcates sequences with a “pass” flag if it considers them most likely to be biologically real transcripts, and TATAT uses this “pass” flag to determine which sequences move forward in the workflow. It is worth mentioning this means that while a different thinning tool could be used, the SQLite database would need to be appropriately updated with a flag to represent which sequences should be used instead. Unless the user has a good understanding of SQL database structure and queries, we encourage users to continue using EvidentialGene and the default code.

#### 2.1.3 Annotation

Working with the knowledge that genes may be evolutionarily conserved, or be recreated through selective pressure, a common approach to annotating unidentified sequences is to perform a BLAST search against previously verified, annotated sequences from different organisms (16,32–35). This method is tried and true and requires little explanation. Likewise, the NCBI’s BLAST tool is consistently used in transcriptomics and is exhaustively validated and supported (20,22,34,36), so we chose to leverage it for annotation.

However, the entire NCBI nucleotide database is very large (∼560 GB at the time of writing) and contains many redundant and intergenic sequences, including entire chromosomes. Additionally, if a researcher knows their samples are bacteria, it may be uninformative and a waste of time to BLAST their bacterial sequences to mammalian genes. Therefore, we do not include a default BLAST database in TATAT and instead provide code and instructions on how to build a BLAST nucleotide database or download the “vertebrata core nt” database we generated for this manuscript (which is nonredundant, genic, and only ∼49 GB) (see 2.1.5).

Once the BLAST search (v2.16.0) is complete, the accession numbers from the subject sequences are collated and submitted in batches to the NCBI servers via their tool Datasets. The NCBI servers return various metadata about the accession numbers, including whether the nucleotide sequence is coding and what its gene symbol is, if any. In the default TATAT workflow, accession numbers that produce coding genes with gene symbols are selected and a table in the SQLite database is generated mapping accession numbers to their gene symbols.

Finally, TATAT generates the “core” coding transcriptome. The process for determining it is as follows: first each sequence’s BLAST hits are examined, and the most statistically significant hit with a true gene symbol, if available, is chosen to annotate the sequence. For example, if a sequence has a BLAST hit for LOC123, followed by UBB, the UBB symbol will be preferentially assigned over LOC123 even if LOC123 is more significant. This facilitates downstream analyses such as Gene Set Enrichment (GSE) which requires unambiguous gene symbols. Then, all the sequences that are annotated with the same gene symbol (e.g. UBB) are clustered, and the longest sequence is chosen to represent the cluster. The logic here is if EvidentialGene and BLAST/Datasets have correctly identified these sequences as real mRNA transcripts, then the longest transcript should be the longest isoform (see Supplemental Figure 1 for reference). This representative sequence is flagged as a “core” gene (the representative isoform), and all the core coding genes may then be written to a fasta file.

By default, TATAT also ignores ambiguous genes when writing the final coding transcriptome file. This is to prevent downstream situations where reads may map to a “LOC” or “CUN” gene, which may siphon all the read counts from a sequence with a real gene symbol (e.g. UBB), but then that ambiguous gene is skipped in subsequent analyses which require a real gene symbol; this would lead to misinterpretation of differential gene expression. However, this behavior can be overridden if desired.

#### 2.1.4 TATAT Benchmarking

To quantify runtime and resource usage we ran the entire TATAT workflow with the same samples and resource requests three times, and report the median runtimes rounded to the nearest 10 minutes. Prior runs during development allowed us to make an educated guess what the maximum RAM usage would be and helped us determine how many CPUs and how much RAM to allocate during benchmarking.

#### 2.1.5 TATAT BLAST Database

In principle, any BLAST database should be compatible with TATAT’s annotation stage. For the analyses performed in this manuscript, we downloaded the NCBI’s “core nt” database on 04/28/2025, then extracted all the vertebrata sequences using the “blastdbcmd” tool and the taxon id 7742. The vertebrata sequences were then built into a BLAST database (version 5) using the “makeblastdb” tool. This final “vertebrata core nt” database is used for annotation and is available on Zenodo (see Data and code availability).

### 2.2 Post TATAT Quality Control

Depending on the novelty of the transcriptome generated by a researcher, follow-up with Quality Control (QC) analysis may be impossible or uninformative. However, our validations are performed on samples from *Rousettus aegyptiacus*, which has a transcriptome on NCBI. Consequently, we directly compare the TATAT generated transcriptome to the NCBI transcriptome.

#### 2.2.1 Gene Symbol Comparison

For the initial analysis we download the *R. aegyptiacus* transcriptome from NCBI (tag GCF_014176215.1). We then generate a set of the gene symbols found from the NCBI dataset and a set from the TATAT transcriptome, remove ambiguous gene symbols (such as LOC), and generate a Venn diagram of the two datasets.

#### 2.2.2 Missing Gene Tissue Expression

The NCBI genes that are not found in the TATAT transcriptome are written to a separate file and utilized for subsequent analysis. Since tissue expression data is not available for *R. aegyptiacus* we instead download the human tissue median TPM expression data from GTEx (tag GTEx_Analysis_v10_RNASeQCv2.4.2; full name provided in the TATAT repository). The expression of each gene per tissue is examined and the tissue with the highest expression is denoted as the gene’s primary tissue. The proportion of missing genes belonging to each primary tissue is calculated and depicted as a pie chart, with the tissues and percentages listed for any above 2.5% of the population. The complete list is available in Supplemental Table 1.

**Table 1.** Benchmarking of the TATAT main workflow. 20 samples totaling 690 GB uncompressed sequencing data were processed through TATAT 3 times. The median runtime was selected and the time per stage reported here, rounded to the nearest 10 minutes. CPUs and RAM listed are requested resources, not the exact amount used (i.e. up to 60 GB could be used, but were not).

| Stage | Runtime | CPUs | RAM | Parallel |
| --- | --- | --- | --- | --- |
| Overall | 8 hr 10 min |  |  |  |
| Assembly | 3 hr 40 min | 10 | 50 GB | 10 jobs |
| Thinning | 1 hr 50 min | 10 | 60 GB | No |
| Annotation | 2 hr 30 min | 20 | 60 GB | No |

#### 2.2.3 Testis Gene Expression

The genes belonging to the testis tissue type are separated by whether or not they are missing from the TATAT transcriptome, then a Mann-Whitney U test is performed, revealing the expression levels of the two groups are statistically different with a p-value = 3.72 × 10^−7^. The groups are also mapped as a violin plot.

#### 2.2.4 Transcript Length Comparison

The transcript lengths for shared genes between the NCBI and TATAT transcriptomes are retrieved and plotted. Only qualitative interpretation of the graph is performed, suggesting most transcripts are roughly the same length.

#### 2.2.5 Transcript Pairwise Alignment

The shared transcripts between NCBI and TATAT undergo pairwise alignment to calculate their sequence identity as a percent. However, often the smaller transcript is used when performing this calculation. Clearly this could give a false representation of how well two sequences align. Consequently, we perform the calculation twice, with either of the sequences as the denominator. Then the proportion of sequence identities in the population are calculated by binning them into 5 groups: 0-20%, 20-40%, 40-60%, 60-80%, 80-100%. An additional calculation is done by removing any transcript pairs where one sequence is less than 80% in length from the other, and the remaining pairs are re-binned into the 5 groups.

#### 2.2.6 Mapping with Salmon

The processed sequencing data for the 20 individual samples are mapped to the TATAT transcriptome using Salmon v1.10.0 (37). First, an indexed version of the transcriptome is generated, then each sample is mapped using default parameters. The exact commands may be found in the TATAT repository.

#### 2.2.7 Multidimensional Scaling Analysis

The sample gene counts generated by Salmon are collated into a single file and used to perform multidimensional scaling (MDS) analysis. All genes with 0 variance are removed from the dataset, as well as genes with extremely low counts (less than 100 summed across all samples). The initial attempt revealed dramatic bias, which appeared to be an artifact from two highly expressed blood-related genes (ALB and HBB; albumin and hemoglobin subunit beta) and a housekeeping gene (EEF1A1; eukaryotic translation elongation factor 1 alpha 1). We reasoned that depending on how the tissues were extracted, the blood genes could be contamination from surrounding vasculature. We decided to exclude these genes and perform the analysis again. The resulting plot clearly shows the samples group by tissue type.

#### 2.2.8 BUSCO Analysis

The CDSs are translated to amino acid sequences with inhouse code and analyzed by BUSCO v5.8.3 (38) with default parameters, using BUSCO’s “vertebrata_odb12” orthologs. The results are compared to other values produced in the initial BUSCO paper. It is also worth noting only enough of BUSCO is installed in TATAT to process amino acid sequences; it cannot handle nucleotide sequences.

### 2.3 Software, Data Analysis, and Visualizations

Most of TATAT relies on publicly available bioinformatic tools, which are described above. It also leverages a SQLite database and several inhouse python scripts. All of these are included in the Docker image.

The post TATAT QC analyses can also be performed using tools and scripts in the Docker image. However, they may not be suitable for all transcriptomes generated.

All graphical visualizations are generated using matplotlib (39) and seaborn (40) in python3, with some minor enhancements to the text labels using Adobe Illustrator. Also Figure 1 and Supplemental Figure 1 are made entirely with Adobe Illustrator.

## 3 Results

To validate TATAT, we wanted to use an externally validated dataset to compare to the TATAT-generate transcriptome. However, we also wanted to test its ability to generate transcriptomes for non-model organisms (e.g. not humans). This led us to select *Rousettus aegyptiacus* for our evaluation dataset, using 20 RNA-seq samples from 11 different tissues described in Lee et al (21). Their metadata is provided in Supplemental Table 2.

### 3.1 Running and Benchmarking TATAT

We processed the 20 samples through the main TATAT workflow (shown in Figure 1). The Assembly stage produced 14,536,735 contigs, which dramatically exceeds the expected number of genes. The Thinning stage reduced this number to 675,117 candidate CDSs (much lower than the 4,746,293 transcripts produced by the thinning method in the Lee et al paper). Finally, the Annotation stage produced an annotated “core” coding transcriptome of 16,766 unambiguous genes.

To evaluate runtime and resource usage, we ran TATAT three times (as described in 2.1.4) and the median values are reported here in Table 1. The entire workflow ran in ∼8 hours, or one workday. Unsurprisingly, the Assembly stage took the longest and, when running 10 assemblies in parallel, required the most resources at 100 CPUs and 500 GB RAM. However, the assemblies could be run sequentially to reduce resource requirements, but this in turn would increase Assembly runtime to about a day on our HPC.

### 3.2 Post TATAT Quality Control Analysis

To validate the TATAT-generated coding transcriptome, we compared it to the *R. aegyptiacus* transcriptome from NCBI. First, the gene symbols present in each transcriptome were examined and it was observed that ∼96% of the named NCBI genes were found in the TATAT transcriptome (Figure 2A). To determine why the remaining 4% were missing, we downloaded human tissue expression values to assign primary tissues of expression to each gene. We were able to determine that most of the missing genes are primarily expressed in tissues not present in the 11 tissues represented in our analysis (Figure 2B, Supplemental Table 1). However, it was noted the single most prevalent tissue for missing genes was testis, which was one of the samples included, albeit with only one replicate. This suggested TATAT may have struggled with low replicate counts, but we explored these genes further by examining the human expression levels between present and missing genes (Figure 2C). We observed that the missing genes overall had lower expression levels (Mann-Whitney U test, p-value = 3.72 × 10^−7^), although some were higher. Collectively, the analyses suggest that TATAT is able to successfully identify most genes (96%), but may struggle when replicate count is low or the transcripts have low expression in the samples processed.

**Figure 2.**
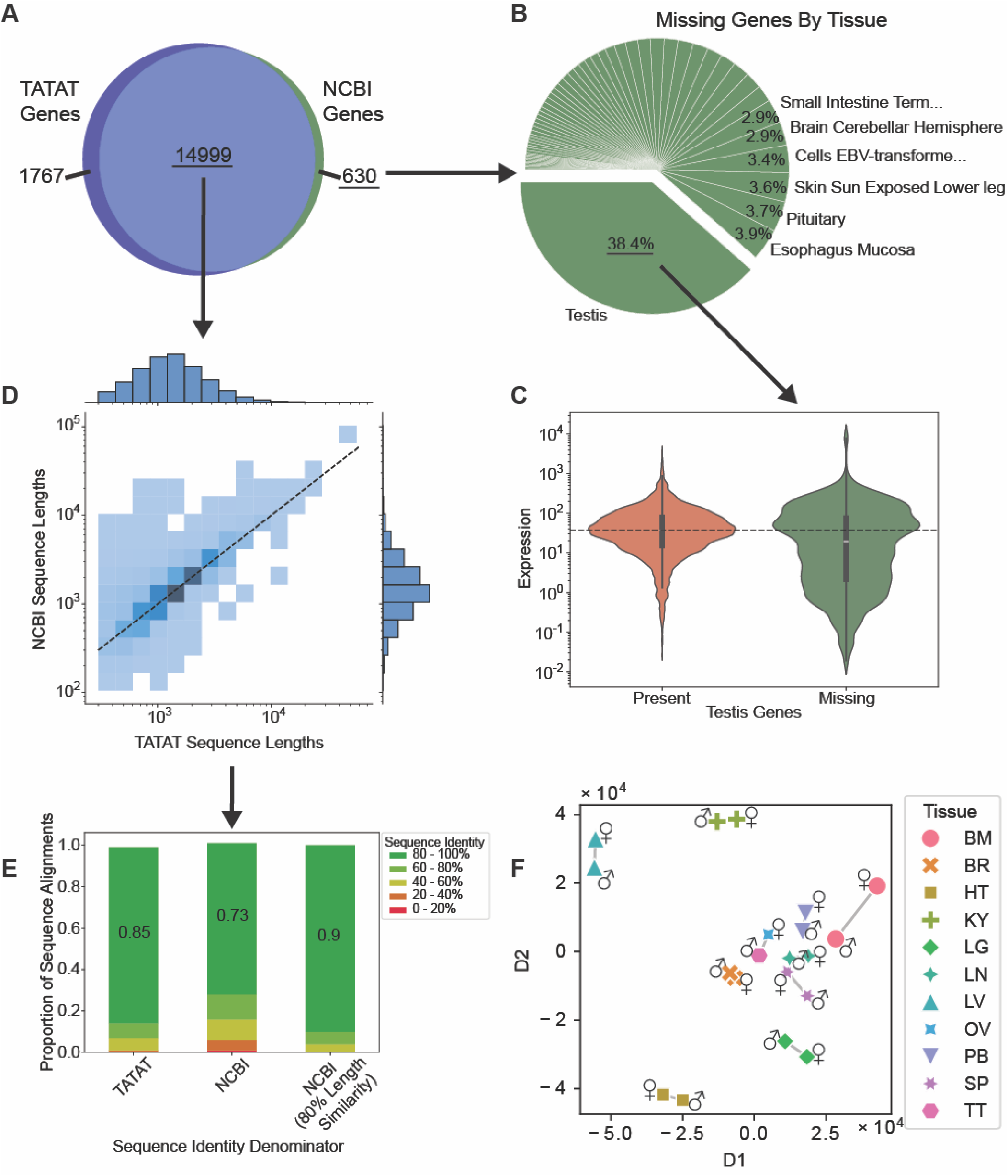
Quality Control Analysis of TATAT-Generated *Rousettus aegyptiacus* transcriptome. (A) Comparison of unambiguous named gene symbols between the TATAT transcriptome and the NCBI transcriptome. ∼96% of NCBI genes were shared with TATAT, but ∼4% were missing. (B) The missing genes were assigned a primary tissue of expression using human expression levels, and the proportion of missing genes per tissue was calculated. Most tissues were not present in the samples, but testis was. (C) The expression levels of the missing testis genes were compared to the expression levels of the present genes. Overall, missing genes had lower expression (Manny-Whitney U test, p-value = 3.72 × 10^−7^). The dashed black line indicates the present genes’ median expression level. (D) All shared genes’ transcript lengths were compared. The dashed black line indicates the 1:1 values and darker boxes indicate higher counts in that bin. Generally, most shared transcript lengths follow the 1:1 line. (E) Sequence identity was calculated per transcript pair, and the final population proportions calculated in bins of 0-20%, 20-40%, 40-60%, 60-80%, 80-100% sequence identity. Numbers in the green bars indicate the proportion of shared transcripts with 80-100% sequence identity. Sequence identities were calculated with either the TATAT or NCBI transcript as the denominator, and using only transcripts with 80% length similarity between the two. (F) The samples’ reads were mapped to the TATAT transcriptome and MDS analysis performed on the counts. Sample types are indicated by shape and color, and sex with gender symbols. The graph shows samples grouped by tissue type.

We then examined the nucleotide sequences between the TATAT and NCBI genes. First a qualitative analysis of the transcript length was done by plotting the lengths in a bivariate histogram (Figure 2D). Manual inspection suggests that the majority of transcripts are of similar length, but the TATAT transcripts tend to be a little shorter. This is not strictly a problem, as shorter transcripts may merely indicate TATAT captured a smaller isoform of the genes with the samples available.

We then performed pairwise alignments between each shared gene. As described in 2.2.5, sequence identities were calculated with either the TATAT or NCBI gene as the denominator and the population of sequences identities converted to proportions (Figure 2E). We observed that most transcripts shared high sequence identity of 80-100%, but that the proportion drops from 0.85 to 0.73 between TATAT to NCBI. Due to the previous observation in sequence length differences, we excluded all transcript pairs that were less than 80% in length between each other, and repeated the calculation, raising the proportion of transcripts with 80-100% sequence identity to 0.90. This suggests that TATAT is able to recreate most genes with the correct sequence and that even low sequence identity may instead be the result of TATAT creating smaller isoforms with the data available.

Following this, we used Salmon to index the TATAT transcriptome and map the reads from the 20 samples to the indexed transcriptome. The gene counts were then used for MDS analysis (2.2.7) and revealed that the samples clustered based on tissue type (Figure 2F). This helps validate that downstream analyses using the TATAT transcriptome would accurately reflect the underlying biology of the samples.

Lastly, we performed BUSCO analysis on the TATAT transcriptome (2.2.8). The results were “C:85.9%[D:0.6%], F:5.5%, M:8.6%” where C represents complete orthologous genes, D is complete and duplicated, F is fragmented genes, and M is missing. The initial BUSCO paper (38) lists the gene sets of model organisms like *Homo sapiens* as producing values of “C:99%[D:1.7%], F:0.0%, M:0.0%” which are clearly better.

However, their Supplemental Table 1 shows that several non-model organisms had lower values, such as *Sus scrofa* (wild boar) which yielded “C:83%[D:7.4%], F:6.8%, M:10%” and *Taeniopygia guttata* (Australian zebra finch) which produced “C:81%[D:3.2%], F:7.5%, M:11%”. Consequently, we believe the TATAT *R. aegyptiacus* values coincide with the expected values of a non-model organism, and these numbers would increase with additional replicates and samples from uncollected tissue types.

## 4 Discussion

In this manuscript we present TATAT, a Dockerized software which contains all its dependencies in a static state to increase the tool’s longevity and to simplifying installation, i.e. downloading a single Docker image. We also preferentially implement tools that are not just accurate, but fast and less resource demanding, making it relatively light weight and able to generate a comprehensive annotated coding transcriptome in a single workday.

Furthermore, we validated the *R. aegyptiacus* transcriptome generated by TATAT by comparing it to the NCBI transcriptome and found a high degree of similarity. Most of the named NCBI genes were recapitulated (∼96%, Figure 2A) with a high degree of nucleotide sequence identity (generally 80-100% identity, Figure 2E). MDS analysis also revealed mapping the original sample reads to the transcriptome produced gene counts which grouped by tissue type (Figure 2F), suggesting downstream analyses would correctly capture information about the underlying biology.

Additionally, while our main objective for validating TATAT was to ensure it could reproduce well curated datasets like those available on NCBI, we also noted TATAT identified 1,767 named genes not present in the current NCBI *Rousettus aegyptiacus* transcriptome. We did not extensively pursue these genes, and therefore can make no definitive statements regarding them. Nonetheless, we tentatively suggest that in addition to the benefits already outlined, TATAT may be able to capture and annotate new genes that other workflows miss.

However, we acknowledge the transcriptomes generated by TATAT contain some limitations. First, TATAT focuses on choosing the longest potential isoform to represent each gene and ignores all shorter potential isoforms in the final transcriptome. This would make it unsuitable for analyses examining isoforms. Second, it appears TATAT may struggle to generate genes when sample replicate count is low (Figure 2B) or expression levels are low (Figure 2C), as suggested by the transcriptome missing some testis genes. These limitations are not unique to TATAT and would likely be overcome with more replicates per tissue type, making it a venial issue. Lastly, TATAT is primarily meant to generate coding transcriptomes, not non-coding. We did include a separate workflow for generating a non-coding RNA (ncRNA) transcriptome (a description is available in the repository), but found it to take significantly longer and more difficult to validate. Therefore, unless a clear use case is identified for a ncRNA transcriptome, we recommend TATAT be used only for coding transcriptome generation.

In summary, TATAT relatively quickly generated a coding transcriptome for *R. aegyptiacus*, which appeared highly accurate and ideal for downstream analyses that rely on changes in gene counts, such as Differential Gene Expression (DGE), Gene Set Enrichment (GSE), and Gene Ontology Enrichment (GOE). However, we would caution against analyses that require extremely high confidence in the nucleotide sequence, such as identifying Single Nucleotide Variants (SNVs), as sequence identity could be improved. Nonetheless, TATAT should expedite transcriptomic analysis for non-model organisms.

## Acknowledgements

This research used resources from the Center for Institutional Research Computing at Washington State University.

## Author contributions

SNS and AJB conceptualized the project and reviewed and edited the manuscript. AJB developed the software, refined the methods, performed validations, and drafted the original manuscript. SNS managed and supervised the project and acquired funding.

## Data and code availability

The code for TATAT is open-source and available under the MIT license at the following GitHub repository: https://github.com/viralemergence/tatat.

The sequencing data for *R. aegyptiacus* was downloaded from the SRA; BioProject accession number PRJNA300284. The vertebrate BLAST database used to annotate the thinned contigs is available through Zenodo: https://zenodo.org/records/15685806.

