## Supplemental data for "TATAT: a containerized software for generating annotated coding transcriptomes from raw RNA-seq data"

**Supplementary data**

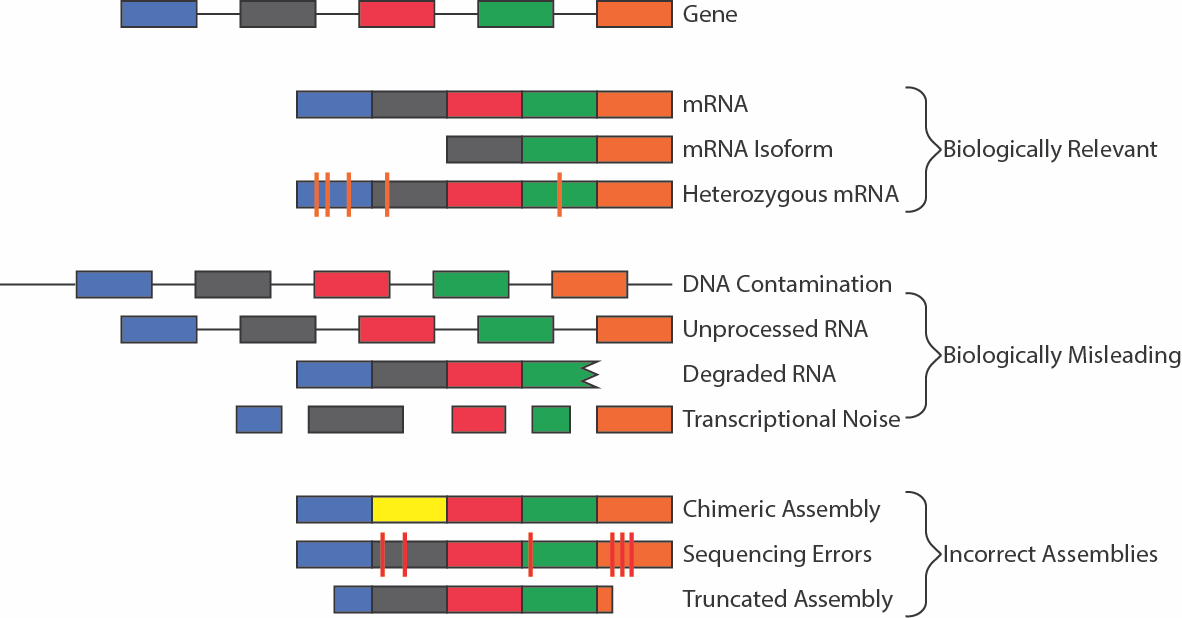

**Supplemental Figure 1 Different sources of contig over-assembly.** This figure shows a gene of interest with different exons colored blue, gray, red, green, and orange, and with introns depicted as black connecting lines. The “Biologically Relevant” assemblies include processed mRNA that no longer has introns, are smaller isoforms, and heterozygous mRNA from genes on other chromosomes in polyploid organisms. The “Biologically Misleading” assemblies are DNA or RNA molecules that may exist, but do not truly represent functional transcribed products. DNA contamination would be chromosomal DNA that makes it to sequencing, unprocessed RNA still contains introns, degraded RNA are molecules that were functional but are being broken down by the cell, and transcriptional noise are nonfunctional products made by misfiring of transcriptional machinery. Lastly, “Incorrect Assemblies” are generated through technical or computational errors. Chimeric assemblies are made by *de novo* assemblers creating a contig that does not exist in the organism, sequencing errors are generated during the sequencing step and persist to assembly, and truncated assemblies are the result of assemblers making the transcript too short, potentially due to low coverage or redundant sequences.
